# Binding mechanism of neutralizing Nanobodies targeting SARS-CoV-2 Spike Glycoprotein

**DOI:** 10.1101/2021.04.23.441186

**Authors:** Mert Golcuk, Aysima Hacisuleyman, Burak Erman, Ahmet Yildiz, Mert Gur

**Affiliations:** Department of Mechanical Engineering, Istanbul Technical University (ITU), Istanbul, Turkey; Institute of Bioengineering, Swiss Federal Institute of Technology (EPFL), Lausanne, Switzerland; Chemical and Biological Engineering Department, Koc University, Istanbul, Turkey; Physics Department, University of California, Berkeley, CA, USA; Department of Molecular and Cellular Biology, University of California, Berkeley, CA, USA

**Keywords:** COVID-19, SARS-CoV-2, nanobodies, molecular dynamics, binding free energy

## Abstract

Severe acute respiratory syndrome coronavirus 2 (SARS-CoV-2) enters human cells upon binding of its spike (S) glycoproteins to ACE2 receptors. Several nanobodies neutralize SARS-CoV-2 infection by binding to the receptor-binding domain (RBD) of S protein, but the underlying mechanism is not well understood. Here, we identified an extended network of pairwise interactions between RBD and nanobodies H11-H4, H11-D4, and Ty1 by performing all-atom molecular dynamics (MD) simulations. Simulations of the nanobody-RBD-ACE2 complex revealed that H11-H4 more strongly binds to RBD without overlapping with ACE2 and triggers dissociation of ACE2 due to electrostatic repulsion. In comparison, Ty1 binding results in dissociation of ACE2 from RBD due to an overlap with the ACE2 binding site, whereas H11-D4 binding does not trigger ACE2 dissociation. Mutations in SARS-CoV-2 501Y.V1 and 501.V2 variants resulted in a negligible effect on RBD-ACE2 binding. However, the 501.V2 variant weakened H11-H4 and H11-D4 binding while strengthening Ty1 binding to RBD. Our simulations indicate that all three nanobodies can neutralize 501Y.V1 while Ty1 is more effective against the 501.V2 variant.

---

Nanobodies are single-domain antibodies that are used in disease diagnosis and as drug carriers. ^1^ Because of their low molecular weight (12-30 kDa) and complexity, they can be mass-produced rapidly at a low cost in bacteria or yeast.^2,3^ Recent studies identified several nanobodies, including H11-H4, H11-D4, Ty1, Nb20, Nb6, and Sb23, ^4-8^ as promising drugs for neutralizing activity against SARS-CoV-2 infection. These nanobodies specifically bind to the receptor-binding domain (RBD) of the spike (S) glycoprotein^4-8^ and blocking its interaction with the human angiotensin-converting enzyme 2 (ACE2).^9,10^ While Ty1^5^, Nb20^6^, Nb6^7^, and Sb23^8^ sterically overlap with the ACE2 binding site^5-8^, H11-H4^4^, and H11-D4^4^ prevent ACE2 binding without an overlap^4^. It remains unclear how these antibodies neutralize S-ACE2 interactions with and without overlap with the ACE2 binding sites.

Recently, two SARS-CoV-2 variants (N501Y ^11^ and N501Y/E484K/K417N) were observed at a fast-growing rate across the globe. While it remains unclear whether current treatments are effective against these variants, nanobodies can be rapidly engineered using recombinant methods in order to cope with the mutagenesis of the S protein. These studies would be greatly aided by an understanding of how existing nanobodies neutralize the wild-type (WT) S protein at the molecular level.

To address these challenges, we performed all-atom MD simulations totaling 27.6 µs in length using the recently-solved structures of the RBD of SARS-CoV-2 S protein in complex with the N-terminal peptidase domain (PD) of human ACE2^12^. Simulations were performed for WT, N501Y, and N501Y/E484K/K417N mutants of RBD in the presence or absence of the nanobodies H11-H4 ^4^, H11-D4 ^4^, and Ty1^5^. We also simulated the detachment of the nanobodies from RBD at low pulling speeds (0.1 Å *ns*^−1^) comparable to high-speed atomic force microscopy (AFM) experiments to estimate the binding strength. These simulations revealed additional interactions between RBD and the nanobodies to those observed in the crystal structures^4,5^. Ty1 overlaps with the ACE2 binding site and more strongly binds to RBD than ACE2. In comparison, H11-H4 and H11-D4 do not overlap with the ACE2 binding site and have similar binding strength to ACE2. Instead of the steric clash, these two nanobodies disrupt S-ACE2 interactions by electrostatic repulsion when they are bound to RBD side by side. Simulations on the N501Y mutant showed that it has minor effects on S-ACE2 and S-nanobody interactions. However, the N501Y/E484K/K417N mutant reduced the binding strength of H11-H4 and H11-D4 while increasing that of Ty1. These results suggest engineering principles to neutralize RBD mutations that take place during the pandemic.

## RESULTS AND DISCUSSION

### The Interaction Network between the RBD and Nanobodies

To model the dynamic interactions of the nanobody-RBD binding interface, we used the co-structure of RBD of the SARS-CoV-2 S protein in complex with H11-H4^4^, H11-D4^4^, and Ty1^5^ (Figure 1). The structure was solvated in a water box that contains physiologically relevant salt (150 mM NaCl) concentration. For each nanobody-RBD complex, two sets of cMD simulations, each of 200 ns in length, were performed to determine the formation of a salt bridge^14^ and a hydrogen bond, as well as electrostatic and hydrophobic interactions between RBD and PD (see Methods).

**Figure 1.**
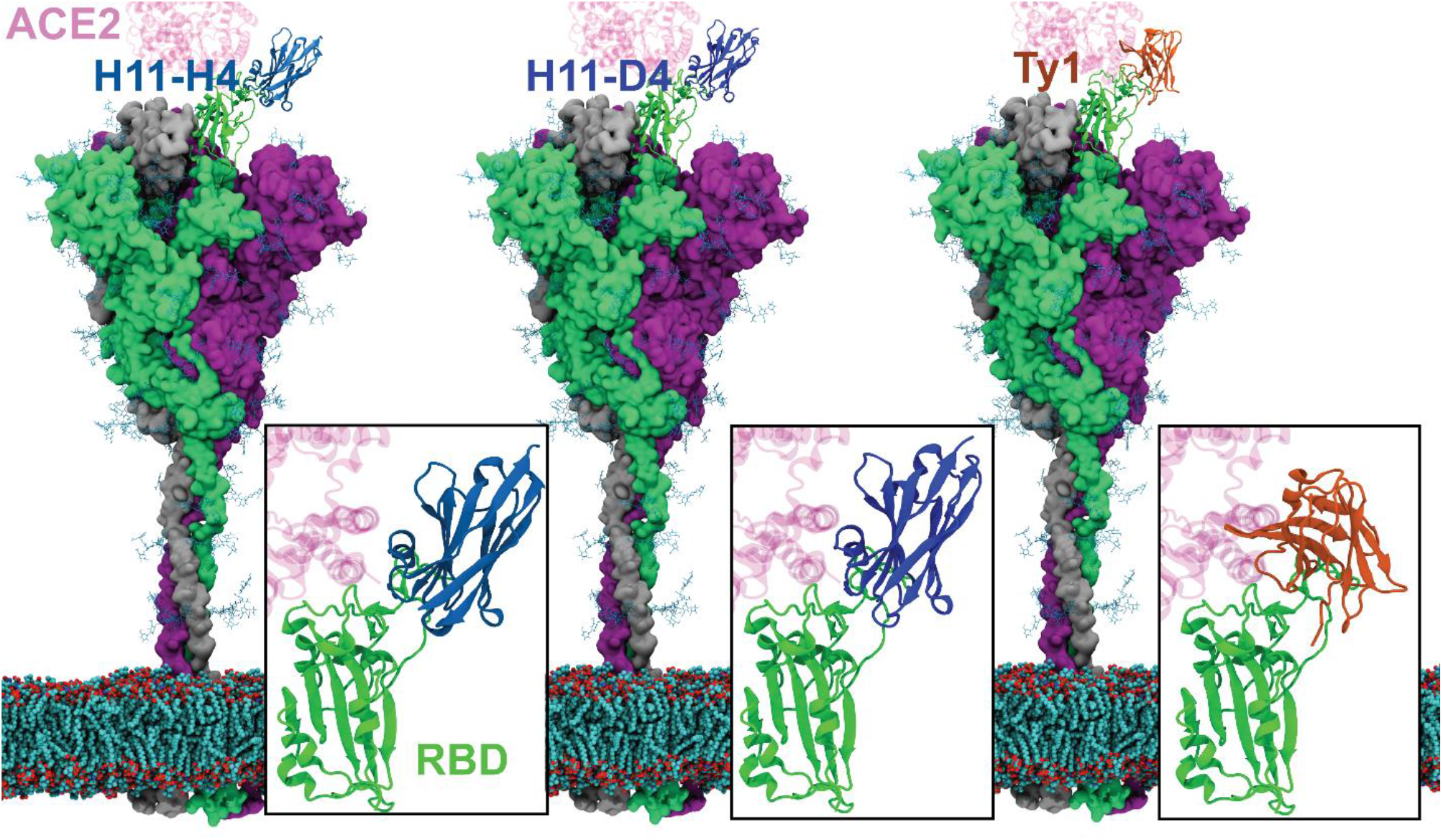
Nanobodies in complex with the RBD of S protein located on the viral membrane. The atomic models were constructed using the full length S protein model^13^, the crystal structure of RBD in complex with ACE2 (PDB: 6M17), and the crystal structures of RBD in complex with the nanobodies H11-H4, H11-D4, and Ty1 (PDB: 6ZBP, 6YZ5, and 6ZXN, respectively). RBD-nanobody interfaces are highlighted in boxes. ACE2 is superimposed on the crystal structures based on its coordinates in the crystal structure (PDB: 6M0J).

Observation frequencies were classified as high and moderate for interactions that occur in 49% and above and between 15-48% of the total trajectory, respectively.^15^

Previously, we divided the RBD-ACE2 interaction surface into three contact regions (CR1-3, Figure 2) and proposed that RBD-ACE2 interaction is primarily stabilized by hydrophobic interactions in CR1. ^15,16^ Similar to ACE2, we observed that nanobodies form many pairwise interactions with RBD. H11-H4 makes 10 hydrophobic interactions, (Figure 2), 5 hydrogen bonds (Figure 2), 1 salt bridge, and 1 electrostatic interaction with RBD (Figure 2). The contact region 1 (CR1) of RBD comprised about half of these interactions. In comparison, CR1 comprised 5 out of 12 interactions (5 hydrophobic interactions, 5 hydrogen bonds, 2 salt bridges, and 2 electrostatic interactions) between H11-D4 and RBD (Figure 2). Ty1 forms a higher number of interactions (24) than H11-H4 and H11-D4. We identified 18 hydrophobic interactions, 6 hydrogen bonds, and 8 electrostatic interactions between Ty1 and RBD (Figure 2). Most of these interaction sites are located in CR1 (Figure 2).

**Figure 2.**
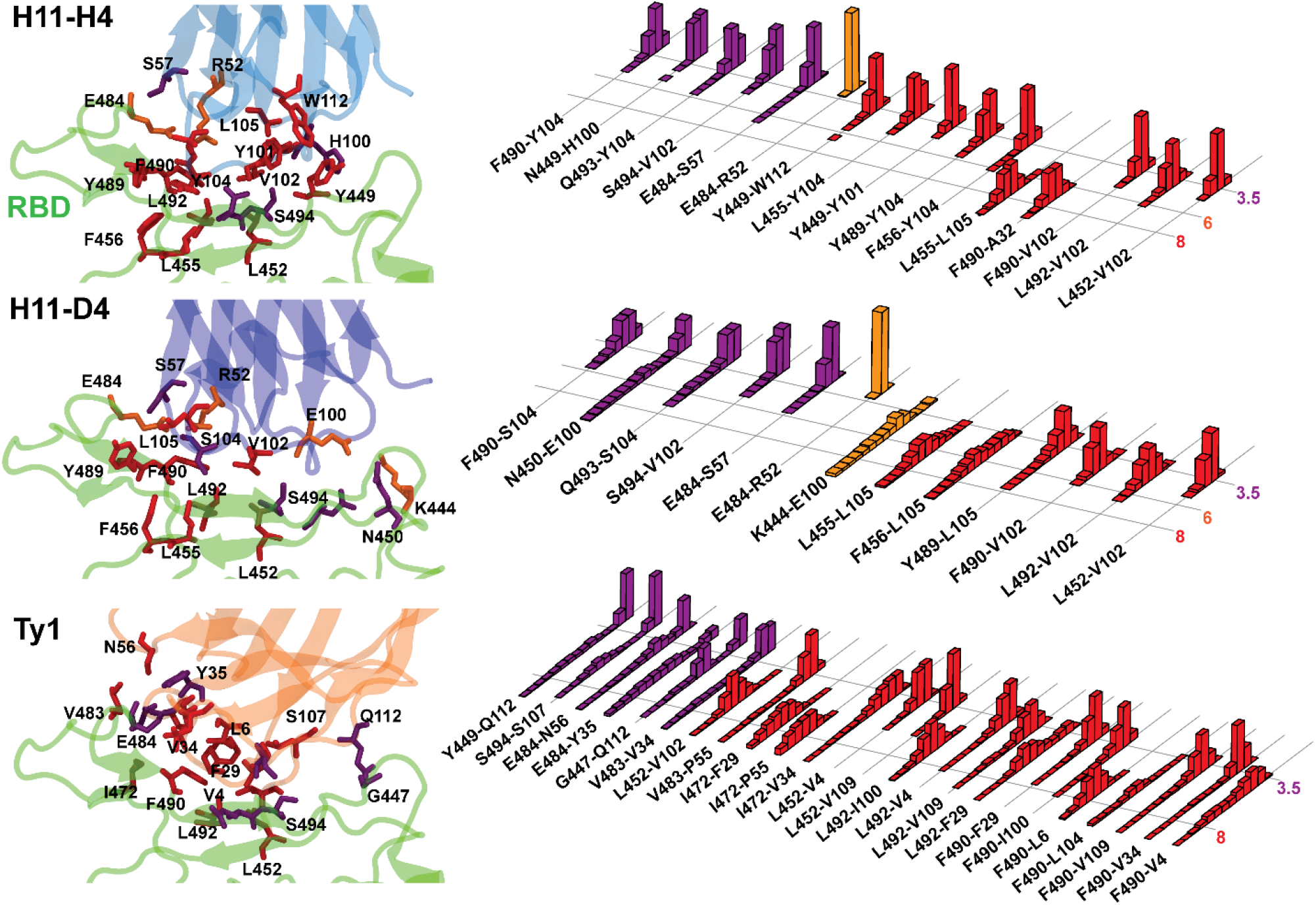
Interactions between RBD of the SARS-CoV-2 S protein and Nanobodies. (Left) Hydrophobic interactions, hydrogen bonding, and salt bridges between (a) H11-H4 (b) H11-D4, and (c) Ty1, with RBD. (Right) Normalized distributions of the distances between the amino-acid pairs that form hydrophobic interactions (red), hydrogen bonds (purple), and salt bridges (orange).

The interaction network we identified between the nanobodies and RBD is mostly consistent with interactions in reported structures. ^4,5^ However, we observed additional 4 hydrophobic interactions (F490-V102, F490-A32, L455-L105, Y449-W112) and 1 hydrogen bond (Q493-Y104) for H11-H4, two hydrophobic interactions (F490-V102, L455-L105), and one salt bridge (K444-E100) for H11-D4, and 6 hydrophobic interactions (L452-V102, L452-V4, L452-V109, L492-F29, L492-V4, L492-I100) and two hydrogen bonds (S494-S107 and G447-Q112) for Ty1. This difference may be due to different thermodynamic conditions between structural studies versus MD simulations, which are performed under physiological conditions.^17^

### ACE2 Release Upon Nanobody Binding

Similar to ACE2, the nanobodies primarily form multiple hydrophobic interactions with CR1 and less strongly with CR2 of RBD. To understand how these nanobodies disrupt S-ACE2 interactions, we first superimposed them together with ACE2 on RBD. We observed that H11-H4 and H11-D4 do not overlap with the ACE2 binding site, whereas Ty1 sterically overlaps with ACE2 (Figure 1). To investigate how H11-H4 and H11-D4 could disrupt S-ACE2 binding without an overlap, we manually docked the structural coordinates of the nanobodies from their co-structure with RBD^4^ onto the RBD-ACE2 complex. We also altered the order of binding by docking the coordinates of ACE2 (PDB ID 6M0J^12^) onto the RBD-nanobody (PDB ID 6ZBP, 6YZ5)^4^ crystal structures (Figure 3). Docking was performed either before or after running conventional MD simulations for 100 ns in solution conditions (Figure 3)

**Figure 3.**
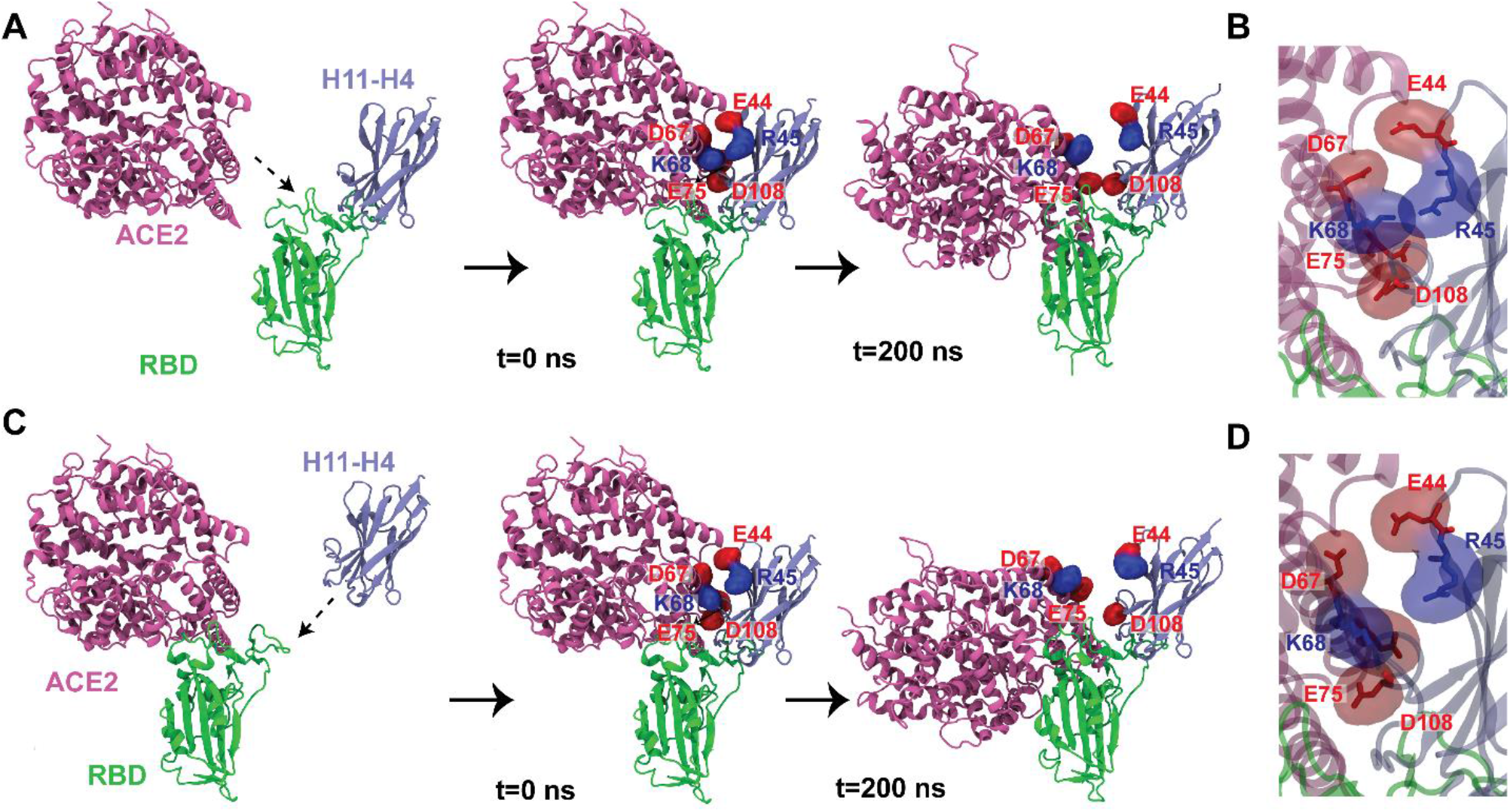
ACE2 is pushed away from RBD by H11-H4 through electrostatic repulsion. A) ACE2 is docked onto the RBD/H11-H4 complex, B) Electrostatic repulsion between ACE2 and H11-H4 upon ACE2 docking. C) H11-H4 is docked onto the RBD/ACE2 complex, D) Electrostatic repulsion between ACE2 and H11-H4 upon H11-H4 docking. Neighboring ACE2 and H11-H4 residues with identical charges are highlighted in surface representation in red (negatively charged) and blue (positively charged). Electrostatic repulsion among these charges dislocates ACE2 from its binding site.

For H11-H4, we ran six 200 ns long conventional MD simulations under different docking orders. In these simulations, H11-H4 binding to RBD abrogated 32.1-90.1% (65% on average) of the high-frequency interactions between RBD and ACE2^15^ (Figure 4, Figure S2). In all cases, ACE2 first detached from CR1, and then lost its binding pose and partially unbound from RBD (Supplementary Video S1). Close examination of the RBD-ACE2-nanobody conformers revealed how nanobody binding results in ACE2 detachment in the absence of direct overlap. We first ruled out the possibility that nanobody binding allosterically alters the interaction surface between ACE2 and RBD. Instead, almost all of the pairwise interactions between RBD and ACE2 remain unaltered, except the E484-K31 interaction was disrupted by the CDR3 loop of H11-H4. We noticed that identically charged residues of ACE2 and nanobodies (D67 with 44, K68 with R45, and E75 with D108) come in close vicinity and repel each other when both proteins are bound to RBD side by side (Figure 3). In all simulations, H11-H4 won the “tug-of-war” and moved ACE2 away from its binding site.

**Figure 4.**
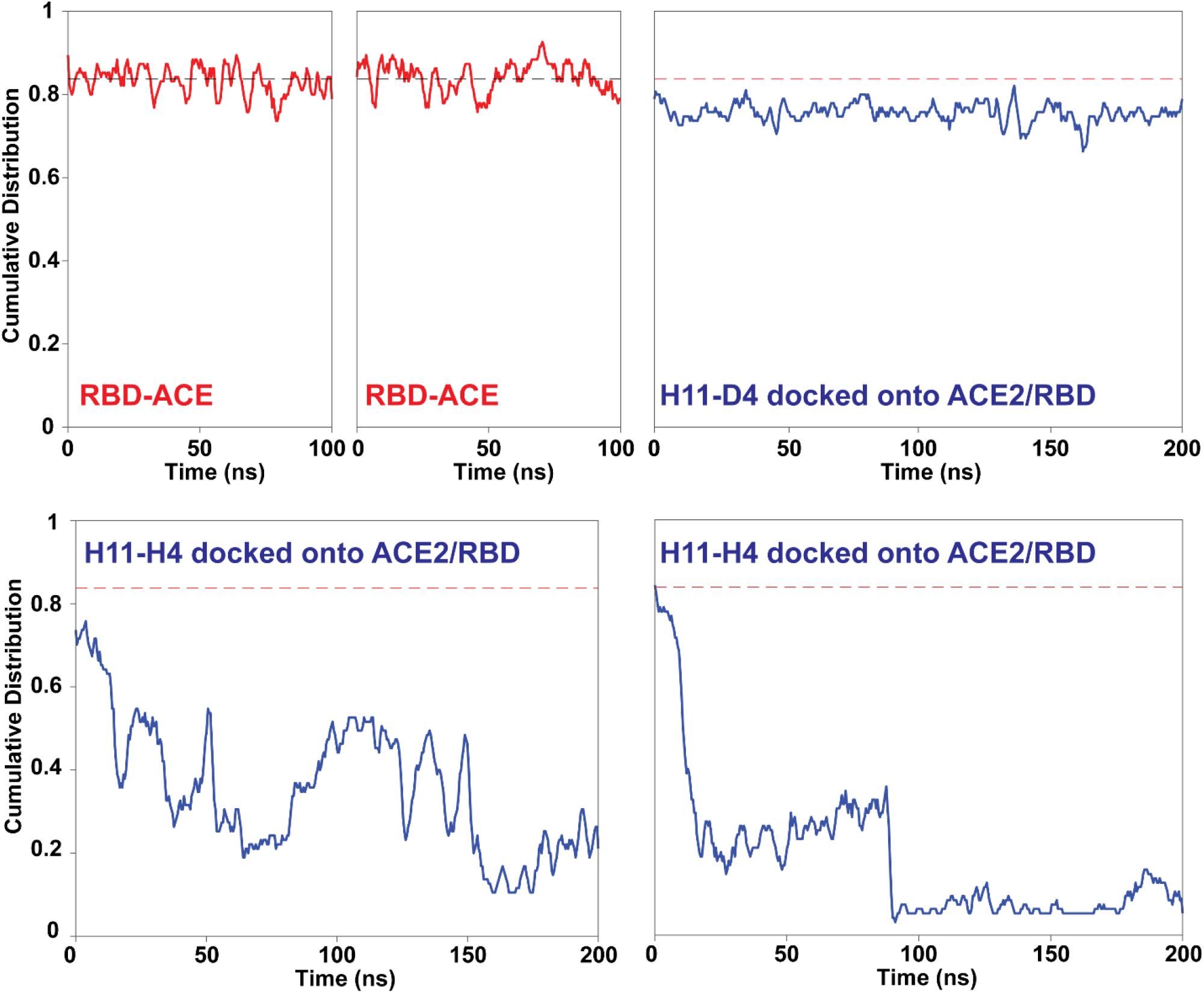
Effect of H11-H4 binding on interactions between RBD and ACE2. Pairwise interactions between RBD and ACE2 were normalized by the total numbers of possible interactions^15^. In the absence of the nanobodies, RBD and ACE2 stably maintained ∼84% of these interactions (red traces). The addition of nanobody to the existing ACE2-RBD complex or the addition of ACE2 to the RBD-nanobody complex resulted in a substantial reduction of pairwise interactions between RBD and ACE2 (blue traces). Time zero indicates the time instant after minimization.

For H11-D4, two 200 ns long MD simulations were performed by docking the nanobody structure on the RBD-ACE2 conformer obtained from the crystal structure (PDB ID 6M0J^12^, 6YZ5^4^). Similar to H11-H4, docking of H11-D4 caused electrostatic repulsion between D67-E44, K68-R45, and E75-D108 (ACE2-nanobody) residues. However, the electrostatic repulsion between ACE2 and nanobody resulted in the disruption of fewer (%62.6 and 34.2%) RBD-ACE2 interactions and ACE2 remained bound to RBD within 200 ns simulation time (Figure 4, Figure S2). These results suggest higher blocking effectiveness of H11-H4 than H11-D4 against ACE2 binding, consistent with higher neutralizing activity H11-H4 against SARS-CoV-2 infection^4^.

### Unbinding of the Nanobodies from S protein under Force

To estimate the binding strength of the nanobodies to RBD, we performed SMD simulations by pulling nanobodies at constant velocities along the vector pointing away from the binding interface (Figure 5). Previous studies typically used 2.5 - 50 Å /ns speeds^18-25^, which overestimate the unbinding free energy. To better estimate the unbinding free energy, we performed our simulations with a velocity of 0.1 Å *ns*^−1^, comparable to the velocities used in high-speed atomic force microscopy (AFM) experiments ^26 15^ Steering forces were applied to the C_α_ atoms of the nanobody residues on the binding interface, whereas C_α_atoms of RBD residues at the binding interface were kept fixed.

**Figure 5.**
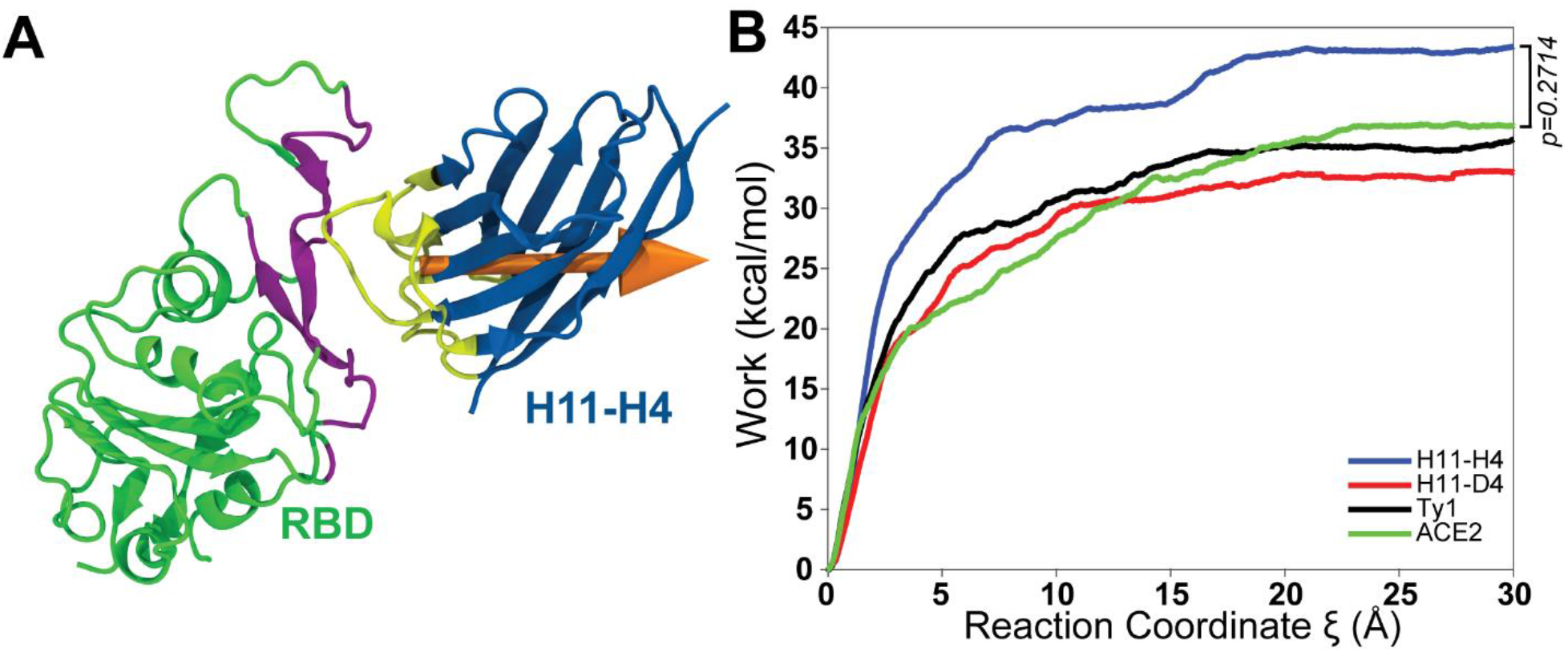
Binding strengths of nanobodies and ACE2 to RBD. (a) In SMD simulations, C_α_ atoms of RBD residues (yellow) were fixed, whereas C_α_ atoms of H11-H4 (purple) were steered at a constant pulling velocity (orange arrow) of 0.1 Å *ns*^−1^. (b) Average work required to move nanobodies along the reaction coordinate. Unbinding work for ACE2 is taken from our previous RBD pulling simulations.^15^

For each nanobody, four 300 ns long SMD simulations (totaling 1200 ns in length, Table S1) were initiated from different conformations sampled from conventional MD simulations. The average work applied to unbind H11-H4, H11-D4, and Ty1 from RBD was 43.4 ± 10.7, 33.0 ± 6.4, and 35.7 ± 6.3 kcal/mol (mean ± s.d.), respectively. These values are comparable to 36.8 ± 9.4 kcal/mol work required to unbind RBD from ACE2 under the same pulling speed^15^. Thus, while ACE2, H11-D4, and Ty1 have similar binding strength to RBD, H11-H4 binds more strongly to RBD. However, based on the Student’s t-test work values were not statistically different.

### Nanobody and ACE2 Interactions of the N501Y and N501Y/E484K/K417N Mutant

Next, we performed conventional and steered MD simulations to investigate how ACE2 and nanobody binding is affected by N501Y (501Y.V1 variant) and N501Y/E484K/K417N (501.V2 variant) mutations on RBD. While N501Y and K417N mutations are located on the ACE2-RBD binding interface, E484K is located on the nanobody-RBD binding interface (Figure 6). The total number of interactions between the nanobodies and 501Y.V1 RBD is similar to that between the nanobodies and WT RBD (Figure S5). However, the number of interactions between H11-H4 and H11-D4 and RBD decreases by 3 and 4, respectively, due to the mutations in 501Y.V2 mutations (Figure S6). Ty1 forms 2 additional hydrophobic interactions and 2 fewer hydrogen bonds with the CR1 of 501Y.V2 RBD compared to WT (Figure S6).

**Figure 6.**
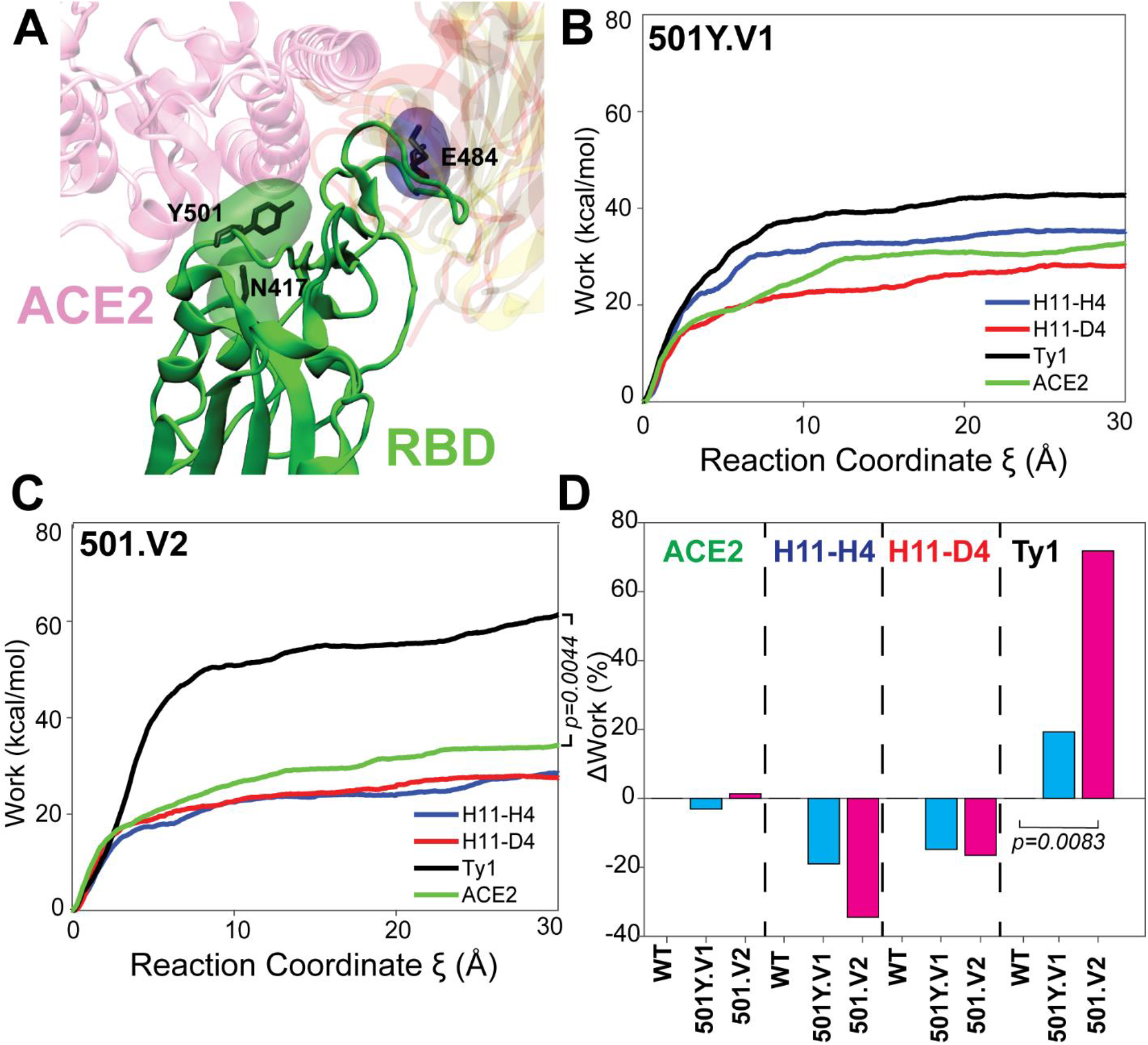
A) Locations of N501Y, E484K, and K417N mutations. Average work required to unbind nanobodies or ACE2 from (B) the N501Y and (C) N501Y/E484K/K417N mutants. D) Relative change of work values due to mutations relative to WT RBD.

We ran SMD simulations to investigate how these mutations affect the RBD binding strength of ACE2 and the nanobodies. We found that ACE2 binds to these mutants as strongly as it binds to WT RBD (Figure 6). Although we observed a slight decrease in the binding strength of H11-H4 and H11-D4 (19% and 15%, respectively) and an increase (19%) in Ty1 binding strength to N501Y, unbinding work values between WT and N501Y were not statistically different (p > 0.13, a two-tailed Student’s t-test). To provide a more extensive sampling of starting conformations, we also ran 20 SMD simulations at higher pulling speeds (2 Å *ns*^−1^). As expected, an increase in pulling speed resulted in higher unbinding work values (Figure S7). Similar to the SMD simulations performed with a pulling speed of 0.1 Å *ns*^−1^, binding strengths of the nanobodies to WT and N501Y RBD were not statistically different under the same velocity and thermodynamic conditions.

Unlike N501Y, the N501Y/E484K/K417N triple mutations decreased the binding strengths of H11-H4 and H11-D4 by 34.5% and 16.5%, respectively, at a pulling speed of 0.1 Å *ns*^−1^. Simulations performed at 2 Å *ns*^−1^ pulling speeds also revealed significantly lower unbinding work values from N501Y/E484K/K417N RBD for both H11-H4 and H11-D4. These changes can be attributed to the loss of a salt bridge and a hydrogen bond due to the E484K mutation. In contrast to H11-H4 and H11-D4, the binding strength of Ty1 increased by 71.8% (Figure 6), nearly twice as strong as the binding strength of other nanobodies and ACE2 to this mutant (p =0.008, a two-tailed Student’s t-test). This may be due to the formation of an extensive network of interactions between the RBD loop (on which K484 is located) and Ty1 (Figure S11). These results suggest that Ty1 can be more effective than H11-H4 and H11-D4 in neutralizing the 501.V2 variant of SARS-CoV-2.

To investigate whether H11-H4 and H11-D4 binding can abrogate ACE2 binding to the N501Y and N501Y/E484K/K417N mutant RBDs, we performed three sets of conventional MD simulations for the ACE2-RBD_mutant_-nanobody trimeric complex (Figure S8). H11-D4 was unable to displace ACE2 from the RBD mutants, similar to that observed in WT RBD. Although H11-H4 was able to unbind ACE2 from WT RBD in all six simulations, it was able to unbind ACE2 from N501Y RBD in only 1 out of 3 simulations and from N501Y/E484K/K417N RBD in 2 out of 3 simulations. These results suggest that H11-H4 is less effective in preventing ACE2 binding to the 501Y and 501Y.V2 variants compared to the WT SARS-CoV-2.

## CONCLUSIONS

We performed an extensive in *silico* analysis to explore how nanobodies H11-H4, H11-D4, and Ty1 disrupt S-ACE2 interactions, and whether these nanobodies can also neutralize the SARS-CoV-2 variants 501Y.V1 and 501.V2. Pulling nanobodies away from WT and mutant S protein RBDs at velocities comparable to those applied at atomic force microscopies enabled us to estimate the nanobody binding strengths and how binding strength is affected by the RBD mutations. Furthermore, by docking of nanobodies (H11-H4 and H11-D4) onto the ACE2-RBD complex or ACE2 onto the RBD-nanobody complexes, we provided a mechanistic explanation of how nanobodies abrogate ACE2 binding from WT and mutant RBD.

Our simulation showed that H11-H4 binds to RBD stronger than H11-D4 and Ty1 and also ACE2. Furthermore, H11-H4 was able to abrogate ACE2 binding in all 6 sets of simulations. On the contrary, H11-D4 binding strength is indistinguishable from ACE2 and it was not able to abrogate ACE2 binding during the length of our simulations. Ty1 also showed similar binding strength as ACE2 to RBD. Taken all together, our MD simulations suggest that H11-H4 is the most effective inhibitor for WT RBD among the three nanobodies investigated in this study.

H11-H4’s effectiveness in abrogating ACE2 binding was diminished by N501Y and N501Y/E484K/K417N mutations of RBD. H11-D4 also did not show abrogation of ACE2 binding within our simulation lengths. These results are consistent with recent studies that reported a minimal and moderate impact of antibodies on 501Y.V1 and 501.V2 variants. ^27-32^ Strikingly, Ty1 binds to the triple RBD mutant two times stronger than ACE2. Super positioning of crystal structures shows that Ty1 sterically blocks ACE2 binding, whereas H11-H4 and H11-D4 did not show direct steric overlap with ACE2. Thus, based on the high binding strength of Ty1 to N501Y/E484K/K417N RBD in comparison to ACE2, we predict that Ty1 will be able to neutralize 501.V2 variants by sterically blocking ACE2 binding.

## METHODS

### MD Simulations System Preparation

As was done in our previous study^15^, for cMD simulations of the S-ACE2 complex the crystal structure of SARS-CoV-2 S protein RBD bound with ACE2 at 2.45 Å resolution (PDB ID: 6M0J)^12^ was used as a template. For the nanobodies crystal structures (PDB ID 6ZBP) ^4^, (PDB ID 6YZ5) ^4^, and (6ZXN)^5^ were used as templates for H11-H4 - RBD, H11-D4 - RBD, and Ty1-RBD systems. The chloride ion, zinc ion, glycans, and water molecules in the crystal structure were kept in their original positions. Each system was solvated in a water box (using the TIP3P water model) having 25 Å cushion in each directions using VMD.^33^ This puts a 50 Å water cushion between the protein complexes and their periodic images. For SMD simulations systems were solvated having 50Å cushion in the positive x directions and 15Å cushion in all other directions, creating enough space for unbinding simulations. Ions were added to neutralize the system and salt concentration was set to 150 mM to construct a physiologically relevant environment. The size of each solvated system was ∼164,000 atoms. All system preparation steps were performed in VMD.^33^

### Conventional MD Simulations

All MD simulations were performed in NAMD 2.13^34^ using the CHARMM36^35^ force field with a time step of 2 fs. MD simulations were performed under N, P, T conditions. The temperature was kept at 310 K using Langevin dynamics with a damping coefficient of 1 ps^-1^. The pressure was maintained at 1 atm using the Langevin Nosé-Hoover method with an oscillation period of 100 fs and a damping time scale of 50 fs. Periodic boundary conditions were applied. 12 Å cutoff distance was used for van der Waals interactions. Long-range electrostatic interactions were calculated using the particle-mesh Ewald method. For each system; first, 10,000 steps of minimization followed by 2 ns of equilibration was performed by keeping the protein fixed. The complete system was minimized for additional 10,000 steps, followed by 4 ns of equilibration by applying constraints on C_α_ atoms. Subsequently, these constraints were released and the system was equilibrated for an additional 4 ns before initiating the production runs. The length of the equilibrium steps is expected to account for the structural differences due to the radically different thermodynamic conditions of crystallization solutions and MD simulations.^17^ MD simulations were performed in Comet and Stampede2 using ∼8 million core-hours in total.

### Criterion for interactions

As a salt bridge formation criteria, a cutoff distance of 6 Å between the basic nitrogens and acidic oxygens was used.^14^ A maximum 3.5 Å distance between hydrogen bond donor and acceptor and a 30° angle between the hydrogen atom, the donor heavy atom, and the acceptor heavy atom was used to score a hydrogen bond formation,.^36^ Interaction pairs that did not satisfy the angle criterion, but the distance criterion were analyzed as electrostatic interactions. A cutoff distance of 8 Å between the side chain carbon atoms was used for hydrophobic interactions.^37-39^

### RMSF Calculations

RMSF values were calculated as 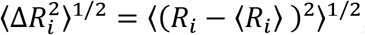, where, ⟨*R*_*i*_⟩ is the mean atomic coordinate of the i^th^ C_α_ atom and *R*_*i*_ is its instantaneous coordinate.

### SMD Simulations

SMD^40^ simulations were used to explore the unbinding process of RBD from ACE2 on time scales accessible to standard simulation lengths. SMD simulations have been applied to explore a wide range of processes, including domain motion,^41,42^ molecule unbinding,^43^ and protein unfolding.^44^ In SMD simulations, a dummy atom is attached to the center of mass of ‘steered’ atoms via a virtual spring and pulled at constant velocity along the ‘pulling direction’, resulting in force *F* to be applied to the SMD atoms along the pulling vector,^34^

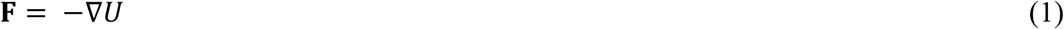

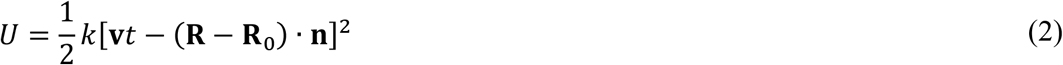

where *U* is the guiding potential, *k* is the spring constant, **v** is the pulling velocity, *t* is time, **R** and **R**_0_ are the coordinates of the center of mass of steered atoms at time *t* and 0, respectively, and **n** is the direction of pulling.^34^ Total work (*W*) performed for each simulation was evaluated by integrating *F* over displacement *ξ* along the pulling direction as = 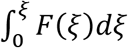.

In SMD simulations of SARS-CoV-2, C_α_ atoms of ACE2 residues S19-S43, T78-P84, Q325-N330, G352-I358, and P389-R393 were kept fixed, whereas C_α_ atoms of RBD residues K417-I418, G446-F456, Y473-A475, and N487-Y505 were steered (Figure 5). Steered atoms were selected as the region comprising the interacting residues. For SARS-CoV SMD simulations the same ACE2 residues were kept fixed. However, two slightly different steered atoms selections were applied: i) Using the same residue positions as for SARS-CoV-2, which are V404-I405, T433-L443, F460-S461, and N473-Y491, and ii) selecting the region comprising the interacting residues, which areT433-L443, F460-D463, and N473-Y491. The total number of fixed and steered atoms were identical in all simulations. The pulling direction was selected as the vector pointing from the center of mass of fixed atoms to the center of mass of steered atoms. The pulling direction also serves as the reaction coordinate ξ for free energy calculations. Each SMD simulation was performed for 300 ns using a 0.1 Å *ns*^−1^ pulling velocity. At a spring constant of 100 *kcal mol*^−1^ Å ^−2^, the center of mass of the steered atoms followed the dummy atom closely while the spring was still soft enough to allow small deviations. For each system, 4 conformations were sampled with a 100 ns frequency (100 and 200 ns time instant). These conformations served as 4 separate starting conformations, **R**_0_, for each set of SMD simulations.

## ASSOCIATED CONTENT

Supplementary Figures S and Table S (PDF).

Supplementary movie S1 (AVI).

Supplementary movie S2 (AVI).

## AUTHOR INFORMATION

## Author

Mert Golcuk – Department of Mechanical Engineering, Istanbul Technical University, Istanbul, 34437, Turkey; https://orcid.org/0000-0001-5476-8160;

Ahmet Yildiz – Department of Physics, University of California Berkeley, 474 Stanley Hall, Berkeley, CA, 94720-3220, USA; https://orcid.org/0000-0003-4792-174X

## Author Contributions

### Funding Sources

This work is supported by COVID-19 HPC Consortium (Grant number: TG-BIO200053)

## ACKNOWLEDGMENT

We gratefully acknowledge the support of the COVID-19 HPC Consortium (Grant number: TG-BIO200053) and Extreme Science and Engineering Discovery Environment (XSEDE)

## ABBREVIATIONS

µs: microsecond
ACE2: angiotensin-converting enzyme 2
atm: standard atmosphere
Cα: carbon alpha
cMD: conventional molecular dynamics
COVID-19: coronavirus disease 2019
CR: contact region
fs: femtosecond
MD: molecular dynamics
NAMD: nanoscale molecular dynamics
ns: nanosecond
PD: peptidase domain
ps: picosecond
RBD: receptor-binding domain
RMSF: root mean square fluctuation
RNA: ribonucleic acid
S: spike
SARS-CoV: severe acute respiratory syndrome-coronavirus
SMD: steered molecular dynamics
VMD: visual molecular dynamics
WT: wild-type

## References

1 Bannas, P., Hambach, J. & Koch-Nolte, F. Nanobodies and nanobody-based human heavy chain antibodies as antitumor therapeutics. Frontiers in immunology 8, 1603 (2017).

2 Gonzalez-Sapienza, G., Rossotti, M. A. & Tabares-da Rosa, S. Single-domain antibodies as versatile affinity reagents for analytical and diagnostic applications. Frontiers in Immunology 8, 977 (2017).

3 Muyldermans, S. A guide to: Generation and design of nanobodies. The FEBS journal (2020).

4 Huo, J. et al. Neutralizing nanobodies bind SARS-CoV-2 spike RBD and block interaction with ACE2. Nature structural & molecular biology 27, 846–854 (2020).

5 Hanke, L. et al. An alpaca nanobody neutralizes SARS-CoV-2 by blocking receptor interaction. Nature communications 11, 1–9 (2020).

6 Xiang, Y. et al. Versatile and multivalent nanobodies efficiently neutralize SARS-CoV-2. Science 370, 1479–1484 (2020).

7 Schoof, M. et al. An ultrapotent synthetic nanobody neutralizes SARS-CoV-2 by stabilizing inactive Spike. Science 370, 1473–1479 (2020).

8 Custódio, T. F. et al. Selection, biophysical and structural analysis of synthetic nanobodies that effectively neutralize SARS-CoV-2. Nature communications 11, 1–11 (2020).

9 Belouzard, S., Millet, J. K., Licitra, B. N. & Whittaker, G. R. Mechanisms of coronavirus cell entry mediated by the viral spike protein. Viruses 4, 1011–1033 (2012).

10 Shang, J. et al. Cell entry mechanisms of SARS-CoV-2. Proceedings of the National Academy of Sciences 117, 11727–11734 (2020).

11 Nuno R. Faria I M.C., Darlan Candido, Lucas A. Moyses Franco, Pamela S. Andrade, Thais M. Coletti, Camila A. M. Silva, Flavia C. Sales, Erika R. Manuli3,4, Renato S. Aguiar, Nelson Gaburo, Cecília da C. Camilo7, Nelson A. Fraiji, Myuki A. Esashika Crispim, Maria do Perpétuo S.S. Carvalho8, Andrew Rambaut, Nick Loman, Oliver G. Pybus, Ester C. Sabino, on behalf of CADDE Genomic Network. Genomic characterisation of an emergent SARS-CoV-2 lineage in Manaus: preliminary findings, <https://virological.org/t/genomic-characterisation-of-an-emergent-sars-cov-2-lineage-in-manaus-preliminary-findings/586> (2021).

12 Lan, J. et al. Structure of the SARS-CoV-2 spike receptor-binding domain bound to the ACE2 receptor. Nature, 1–9 (2020).

13 Casalino, L. et al. Beyond Shielding: The Roles of Glycans in the SARS-CoV-2 Spike Protein. ACS Central Science 6, 1722-1734, doi:10.1021/acscentsci.0c01056 (2020).

14 Beckstein, O., Denning, E. J., Perilla, J. R. & Woolf, T. B. Zipping and unzipping of adenylate kinase: atomistic insights into the ensemble of open↔ closed transitions. Journal of molecular biology 394, 160–176 (2009).

15 Taka, E. et al. Critical Interactions Between the SARS-CoV-2 Spike Glycoprotein and the Human ACE2 Receptor. bioRxiv (2020).

16 Wang, Y., Liu, M. & Gao, J. Enhanced receptor binding of SARS-CoV-2 through networks of hydrogen-bonding and hydrophobic interactions. Proceedings of the National Academy of Sciences (2020).

17 Pullara, F., Wenzhi, M. & GÜR, M. Why protein conformers in molecular dynamics simulations differ from their crystal structures: a thermodynamic insight. Turkish Journal of Chemistry 43, 394–403 (2019).

18 Huang, S. et al. SMD-based interaction-energy fingerprints can predict accurately the dissociation rate constants of HIV-1 protease inhibitors. Journal of chemical information and modeling 59, 159–169 (2018).

19 Moore, D. S., Dalton, J. P. & Tikhonova, I. G. Steered Molecular Dynamic Simulations Reveal Critical Residues for (Un) Binding of Substrates, Inhibitors and a Product of the Malarial PFM1AAP. Biophysical Journal 112, 354a (2017).

20 Yu, Z. et al. Insights from molecular dynamics simulations and steered molecular dynamics simulations to exploit new trends of the interaction between HIF-1α and p300. Journal of Biomolecular Structure and Dynamics 38, 1–12 (2020).

21 Xiao, B.-L. et al. Steered molecular dynamic simulations of conformational lock of Cu, Zn-superoxide dismutase. Scientific reports 9, 1–10 (2019).

22 Célerse, F., Lagardère, L., Derat, E. & Piquemal, J.-P. Massively parallel implementation of Steered Molecular Dynamics in Tinker-HP: comparisons of polarizable and non-polarizable simulations of realistic systems. Journal of chemical theory and computation 15, 3694–3709 (2019).

23 Hu, X. et al. Steered molecular dynamics for studying ligand unbinding of ecdysone receptor. Journal of Biomolecular Structure and Dynamics 36, 3819–3828 (2018).

24 Kosztin, D., Izrailev, S. & Schulten, K. Unbinding of retinoic acid from its receptor studied by steered molecular dynamics. Biophysical journal 76, 188–197 (1999).

25 Azadi, S., Tafazzoli-Shadpour, M. & Omidvar, R. Steered Molecular Dynamics Simulation Study of Quantified Effects of Point Mutation Induced by Breast Cancer on Mechanical Behavior of E-Cadherin. Molecular Biology 52, 723–731 (2018).

26 Rico, F., Gonzalez, L., Casuso, I., Puig-Vidal, M. & Scheuring, S. High-speed force spectroscopy unfolds titin at the velocity of molecular dynamics simulations. Science 342, 741–743 (2013).

27 Wang, P. et al. Antibody resistance of SARS-CoV-2 variants B. 1.351 and B. 1.1. 7. Nature, 1–6 (2021).

28 Shen, X. et al. SARS-CoV-2 variant B. 1.1. 7 is susceptible to neutralizing antibodies elicited by ancestral Spike vaccines. Cell host & microbe (2021).

29 Edara, V. V. et al. Infection and mRNA-1273 vaccine antibodies neutralize SARS-CoV-2 UK variant. medRxiv (2021).

30 Collier, D. A. et al. SARS-CoV-2 B. 1.1. 7 sensitivity to mRNA vaccine-elicited, convalescent and monoclonal antibodies. medRxiv (2021).

31 Wu, K. et al. mRNA-1273 vaccine induces neutralizing antibodies against spike mutants from global SARS-CoV-2 variants. BioRxiv (2021).

32 Emary, K. R. et al. Efficacy of ChAdOx1 nCoV-19 (AZD1222) vaccine against SARS-CoV-2 variant of concern 202012/01 (B. 1.1. 7): an exploratory analysis of a randomised controlled trial. The Lancet (2021).

33 Humphrey, W., Dalke, A. & Schulten, K. VMD: visual molecular dynamics. Journal of molecular graphics 14, 33–38 (1996).

34 Phillips, J. C. et al. Scalable molecular dynamics with NAMD. Journal of computational chemistry 26, 1781–1802 (2005).

35 Best, R. B. et al. Optimization of the additive CHARMM all-atom protein force field targeting improved sampling of the backbone ϕ, ψ and side-chain χ1 and χ2 dihedral angles. Journal of chemical theory and computation 8, 3257–3273 (2012).

36 Durrant, J. D. & McCammon, J. A. HBonanza: a computer algorithm for molecular-dynamics-trajectory hydrogen-bond analysis. Journal of Molecular Graphics and Modelling 31, 5–9 (2011).

37 Stock, P., Utzig, T. & Valtiner, M. Direct and quantitative AFM measurements of the concentration and temperature dependence of the hydrophobic force law at nanoscopic contacts. Journal of colloid and interface science 446, 244–251 (2015).

38 Manavalan, P. & Ponnuswamy, P. A study of the preferred environment of amino acid residues in globular proteins. Archives of biochemistry and biophysics 184, 476–487 (1977).

39 Stavrakoudis, A., Tsoulos, I. G., Shenkarev, Z. O. & Ovchinnikova, T. V. Molecular Dynamics Simulation of Antimicrobial Peptide Arenicin-2: b-Hairpin Stabilization by Noncovalent Interactions.

40 Isralewitz, B., Gao, M. & Schulten, K. Steered molecular dynamics and mechanical functions of proteins. Current opinion in structural biology 11, 224–230 (2001).

41 Izrailev, S., Crofts, A. R., Berry, E. A. & Schulten, K. Steered molecular dynamics simulation of the Rieske subunit motion in the cytochrome bc1 complex. Biophysical journal 77, 1753–1768 (1999).

42 Gur, M. et al. Conformational transition of SARS-CoV-2 spike glycoprotein between its closed and open states. The Journal of Chemical Physics 153, 075101 (2020).

43 Eskici, G. & Gur, M. Computational design of new peptide inhibitors for amyloid beta (Aβ) aggregation in Alzheimer’s disease: application of a novel methodology. Plos one 8 (2013).

44 Lu, H., Isralewitz, B., Krammer, A., Vogel, V. & Schulten, K. Unfolding of titin immunoglobulin domains by steered molecular dynamics simulation. Biophysical journal 75, 662–671 (1998).

