## Supplementary Materials for "Binding mechanism of neutralizing Nanobodies targeting SARS-CoV-2 Spike Glycoprotein"

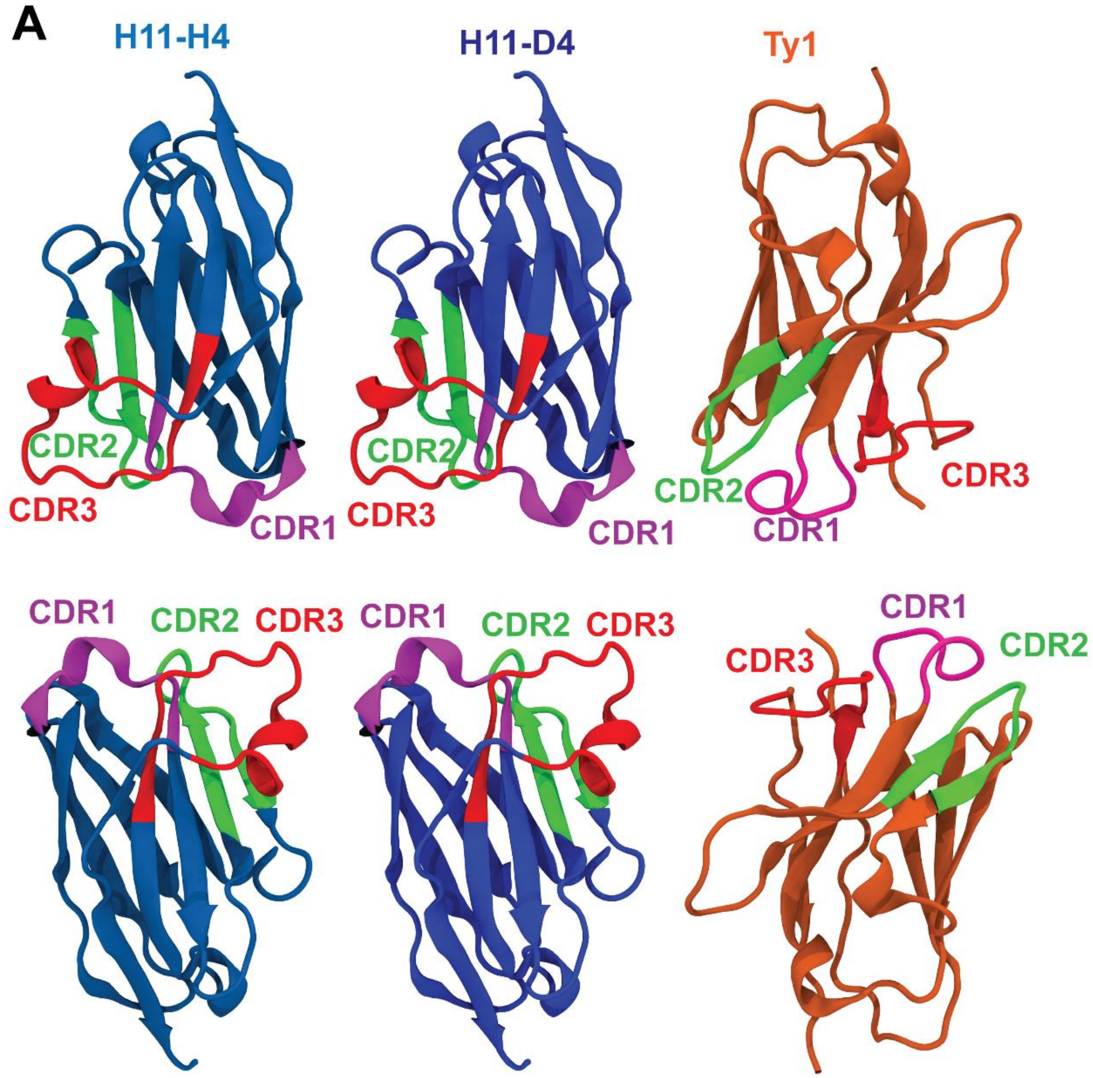


**B**

| | |
|---|---|
| *H11-H4* | Q V Q L V E S G G G L M Q A G G S L R L S C A V S G R T F S T A A M G W F R Q A P G K E R E F V A A I R W S G G S A Y Y A D S V K G R F |
| *H11-D4* | Q V Q L V E S G G G L M Q A G G S L R L S C A V S G R T F S T A A M G W F R Q A P G K E R E F V A A I R W S G G S A Y Y A D S V K G R F |
| *Ty1* | M A Q V Q L V E T G G G L V Q P G G S L R L S C A A S G F T F S S V Y M N W V R Q A P G K G P E W V S R I S P N S G N I G Y T D S V K G |
| | T I S R D K A K N T V Y L Q M N S L K Y E D T A V Y Y C A Q T H Y V S Y L L S D Y A T W P Y D Y W G Q G T Q V T V S S K H H H H H H - - - - - |
| | T I S R D K A K N T V Y L Q M N S L K Y E D T A V Y Y C A R T E N V R S L L S D Y A T W P Y D Y W G Q G T Q V T V S S K H H H H H H - - - - - |
| | R F T I S R D N A K N T L Y L Q M N N L K P E D T A L Y Y C A I G L N L S S S S V R G Q G T Q V T V S S G G L P E T G G H H H H H H - - - - - - |

**Figure S1. (a)** Nanobody structures of H11-H4, H11-D4, and Ty1 provide shown for two different views (top panel and lower panel). CDR1, 2, and 3 are shown in magenta, green, and red, respectively. (b) Sequence alignment of nanobodies.

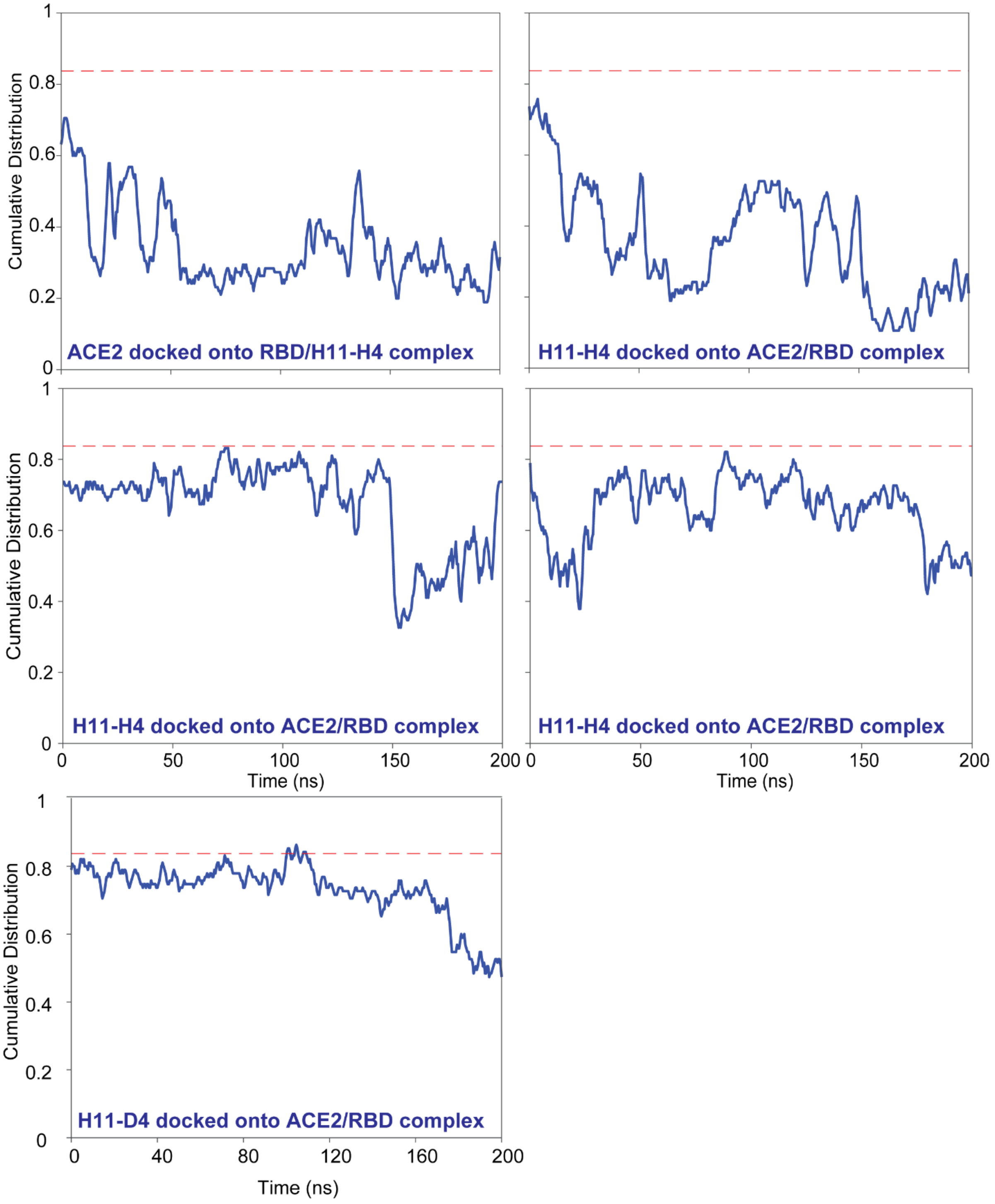


**Figure S2.** Effect of H11-H4 and H11-D4 binding on interactions between ACE2 and S protein. Blue lines indicate the simulations for the ACE2/S/H11-H4 and ACE2/S/H11-D4 trimeric complexes, whereas dashed red lines show the average observed numbers of interactions of the ACE2/S dimeric complex. Time zero indicates the time instant after minimization.

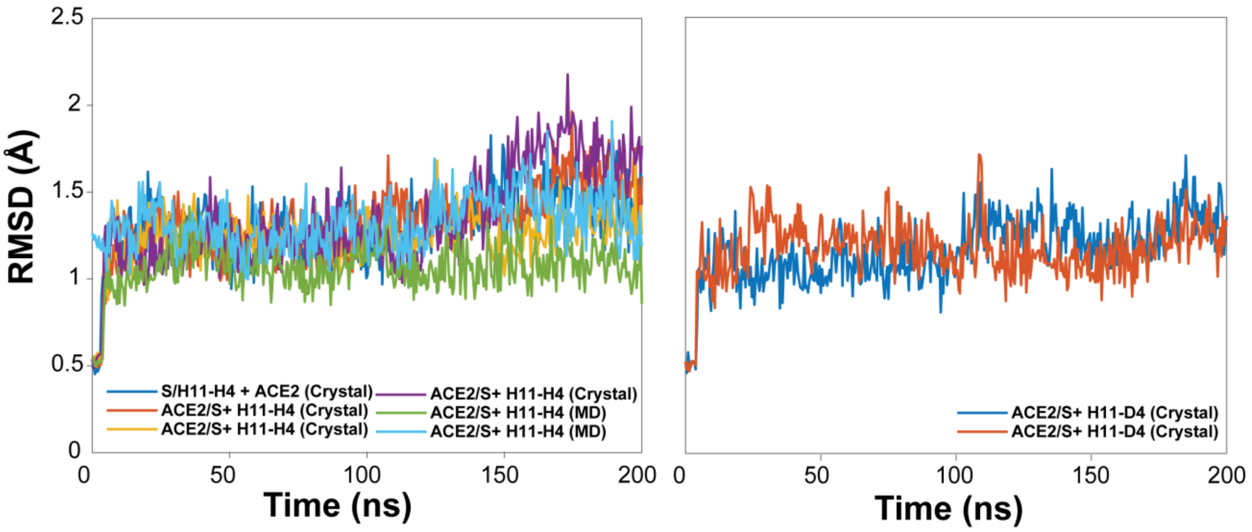


**Figure S3.** Root-mean-square deviation (RMSD) changes of RBD upon nanobody binding in the presence of ACE2.

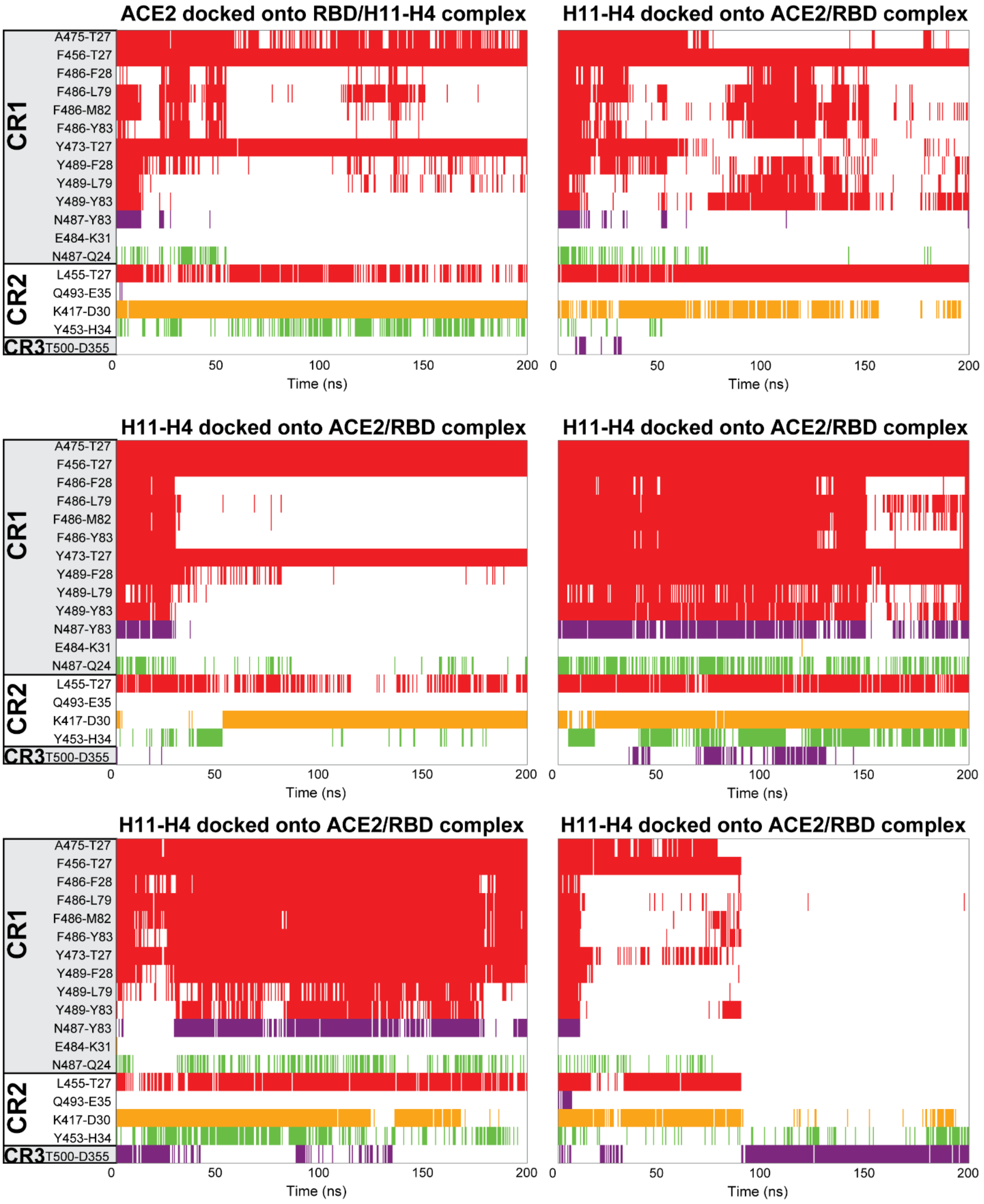


**Figure S4.** Effect of H11-H4 binding on hydrophobic interactions (red), hydrogen bonding (purple), electrostatic interactions (green), and salt bridges (orange) between RBD and ACE2**.** Time zero indicates the time instant after minimization.

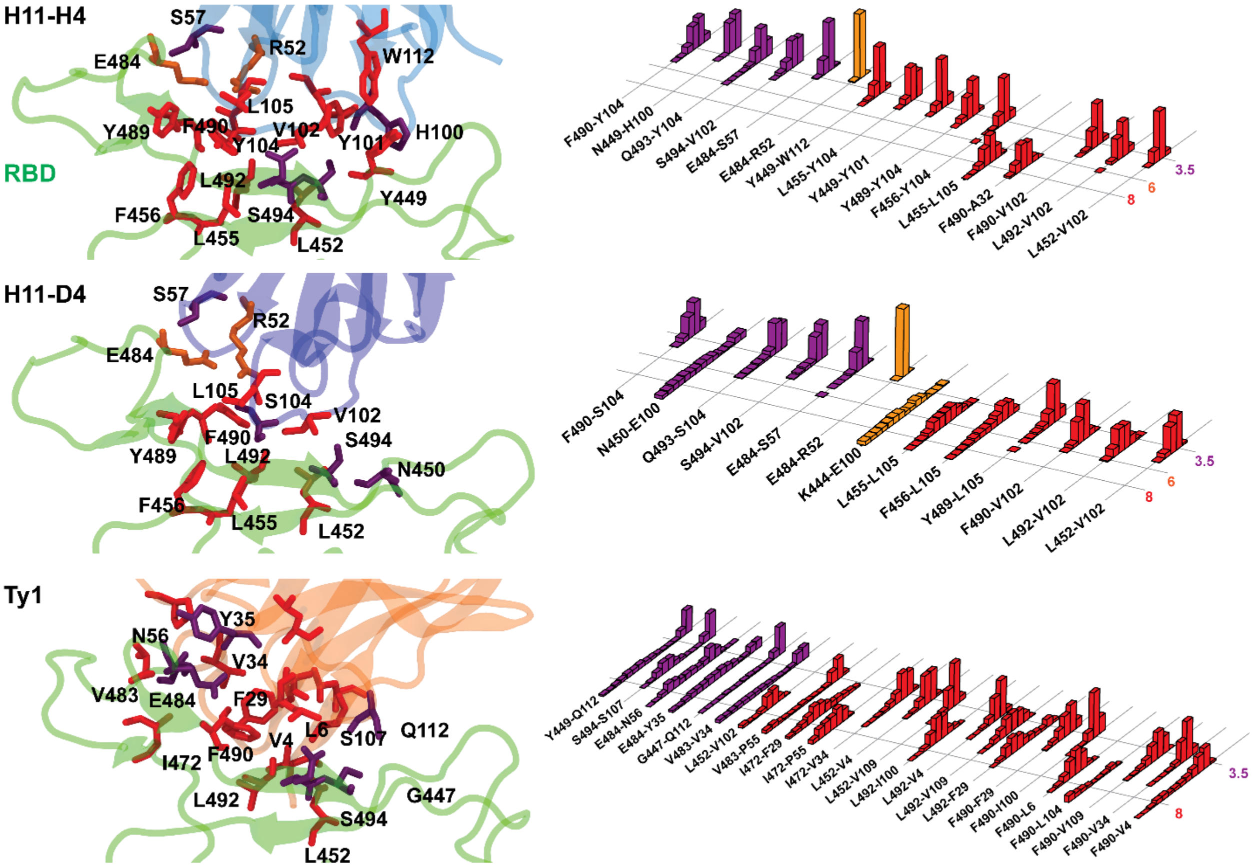


**Figure S5. Interactions between the N501Y mutant of RBD and the nanobodies.** (Left) Hydrophobic interactions, hydrogen bonding, and salt bridges between RBD and (a) H11-H4 (b) H11-D4, or (c) Ty1. (Right) Normalized distributions of the distances between the amino-acid pairs that form hydrophobic interactions (red), hydrogen bonds (purple), and salt bridges (orange).

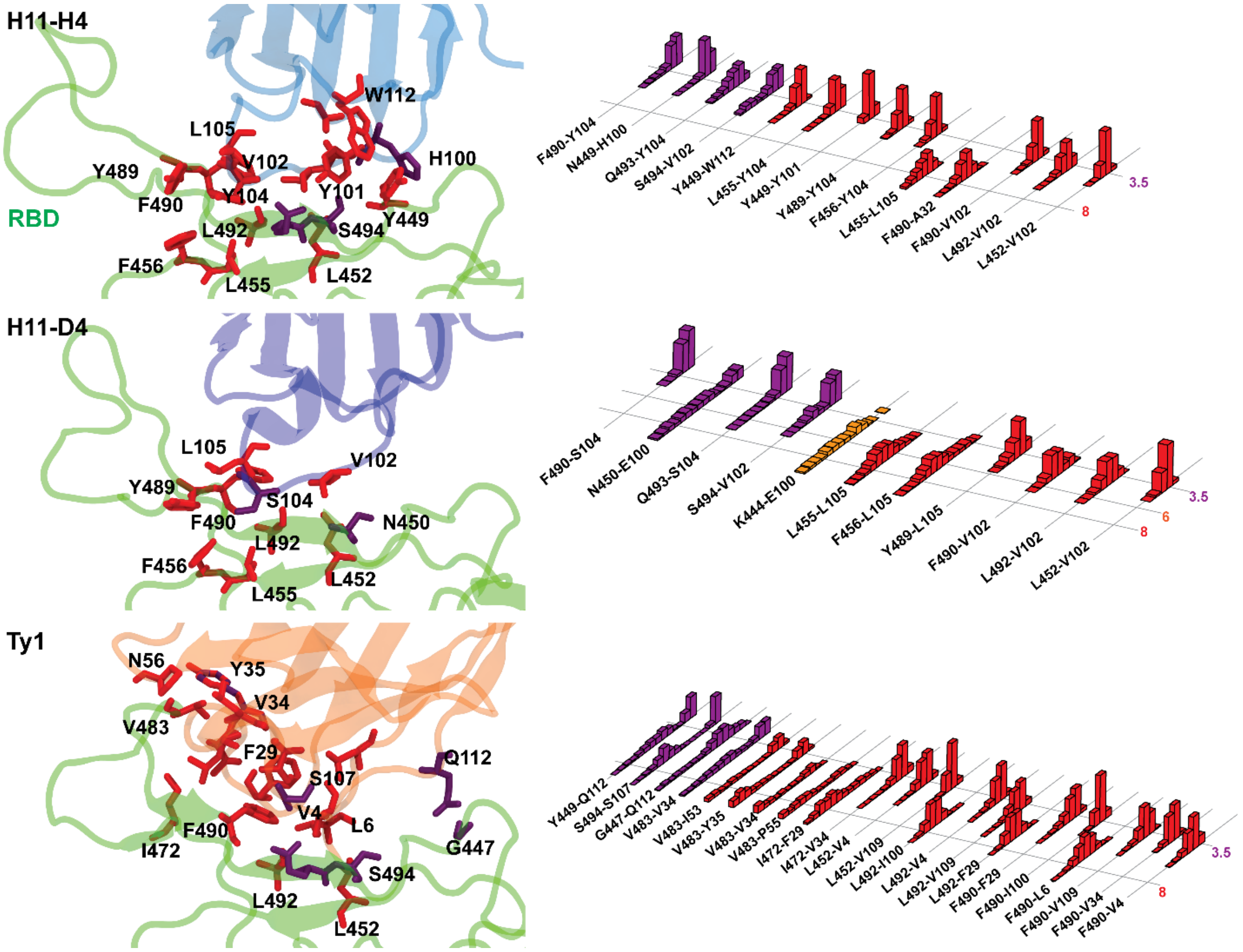


**Figure S6. Interactions between the N501Y/E484K/K417N mutant of RBD and the nanobodies.** (Left) Hydrophobic interactions, hydrogen bonding, and salt bridges between RBD and (a) H11-H4 (b) H11-D4, or (c) Ty1. (Right) Normalized distributions of the distances between the amino-acid pairs that form hydrophobic interactions (red), hydrogen bonds (purple), and salt bridges (orange).

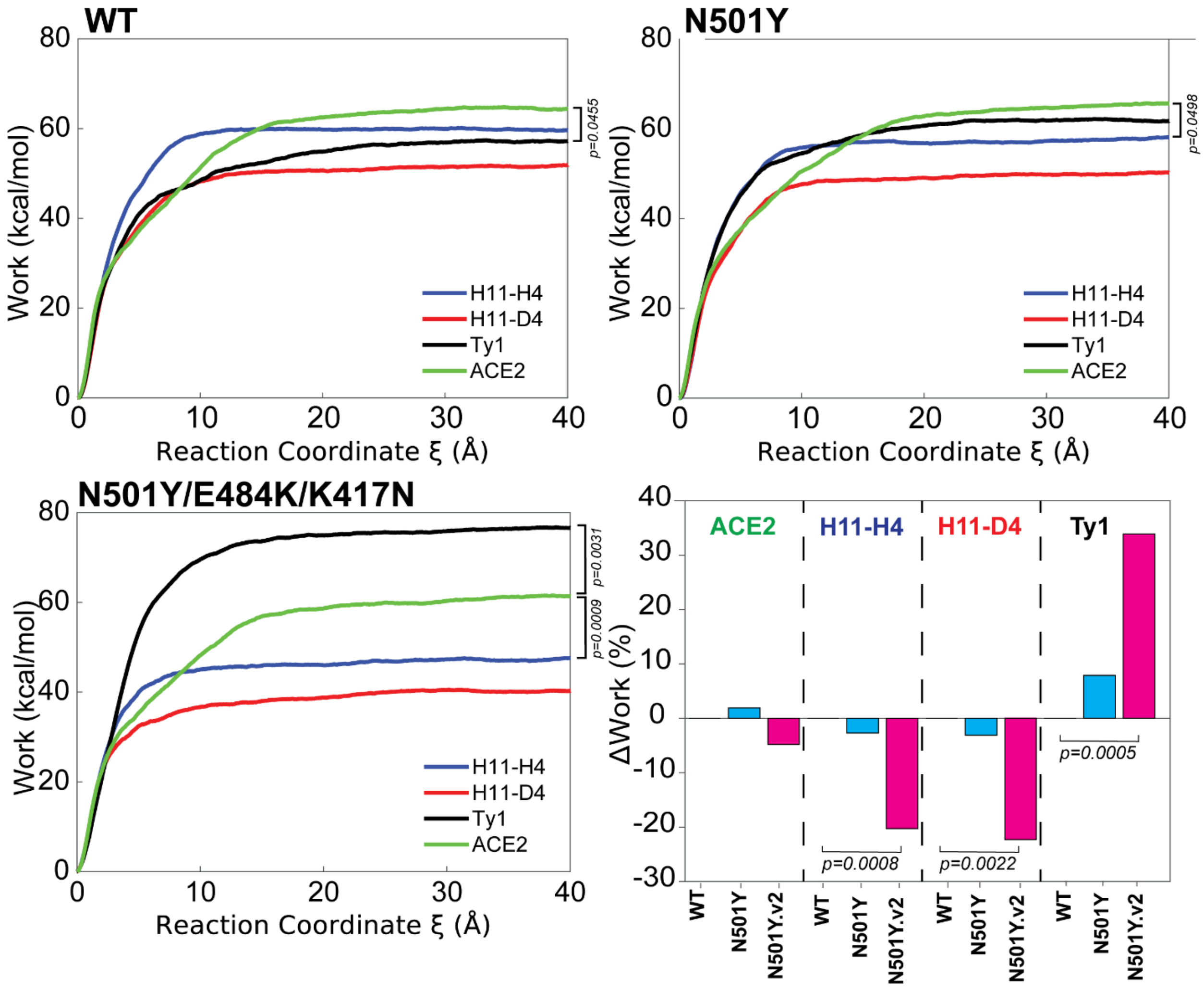


**Figure S7.** Average work required to unbind nanobodies or ACE2 from (a) WT, (b) N501Y, and (C) N501Y/E484K/K417N RBD with 2 Å $ns^{-1}$ pulling speed. D) Relative changes of unbinding work values of RBD mutants compared to WT. p-values are calculated by a two-tailed t-test.

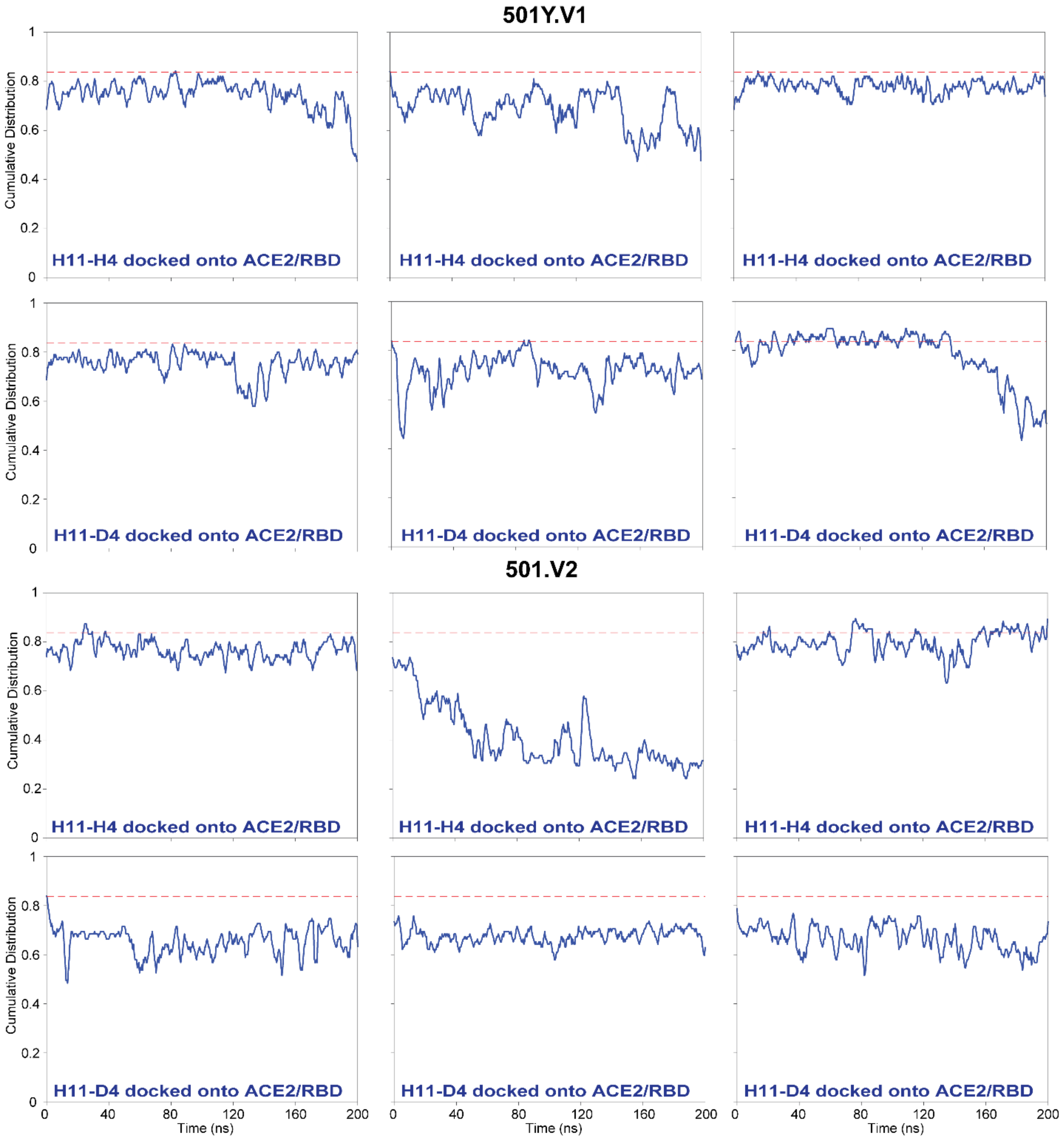


**Figure S8.** Effect of H11-H4 and H11-D4 binding on interactions between ACE2 and S protein variants. Blue lines indicate the simulations for the ACE2/S variant/H11-H4 and H11-D4 trimeric complex, whereas red lines show the simulation results for the ACE2/S dimeric complex. Time zero indicates the time instant after minimization.

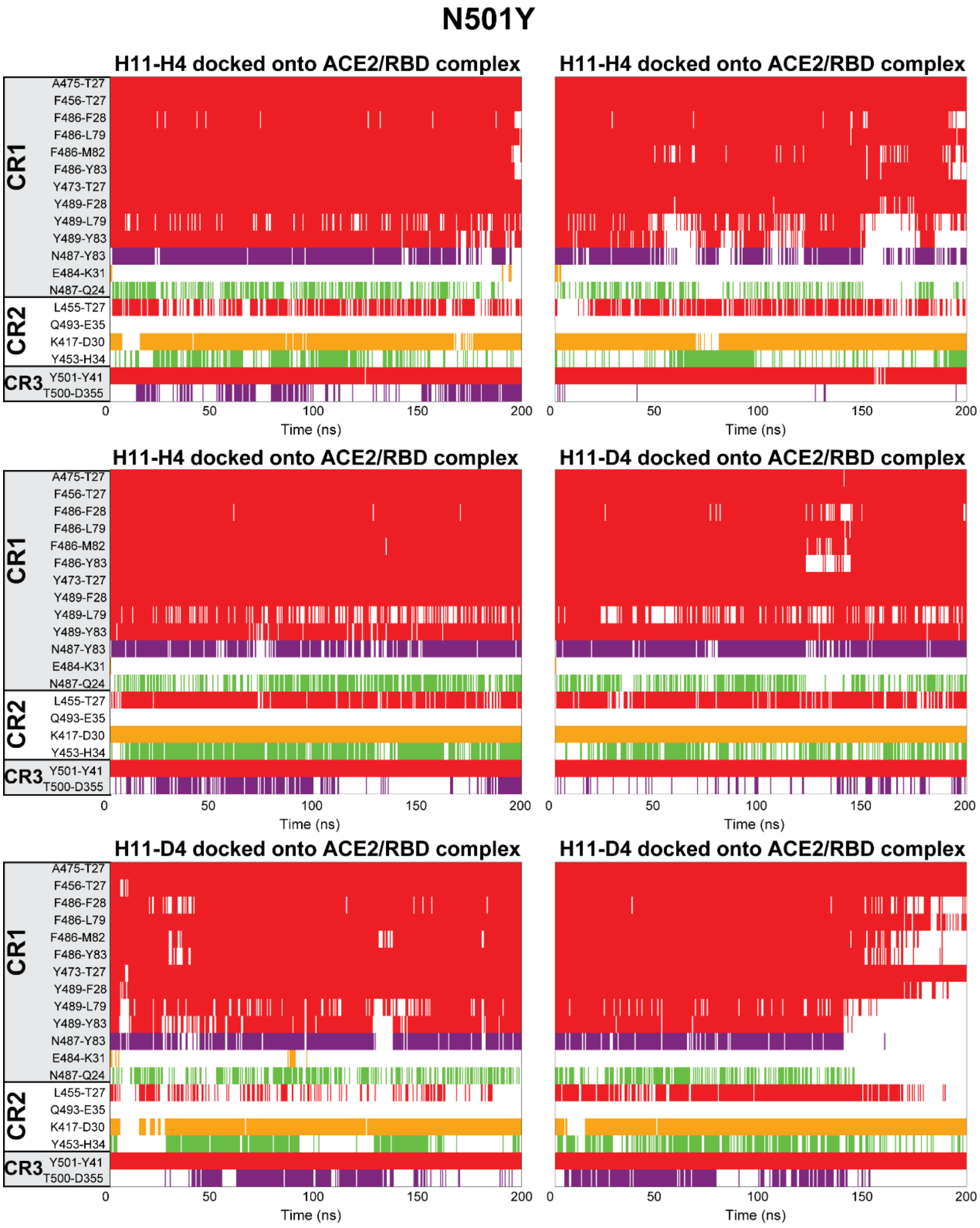


**Figure S9.** Effect of H11-H4 binding on hydrophobic interactions (red), hydrogen bonding (purple), electrostatic interactions (green), and salt bridges (orange) between N501Y mutant of RBD and ACE2**.** Time zero indicates the time instant after minimization.

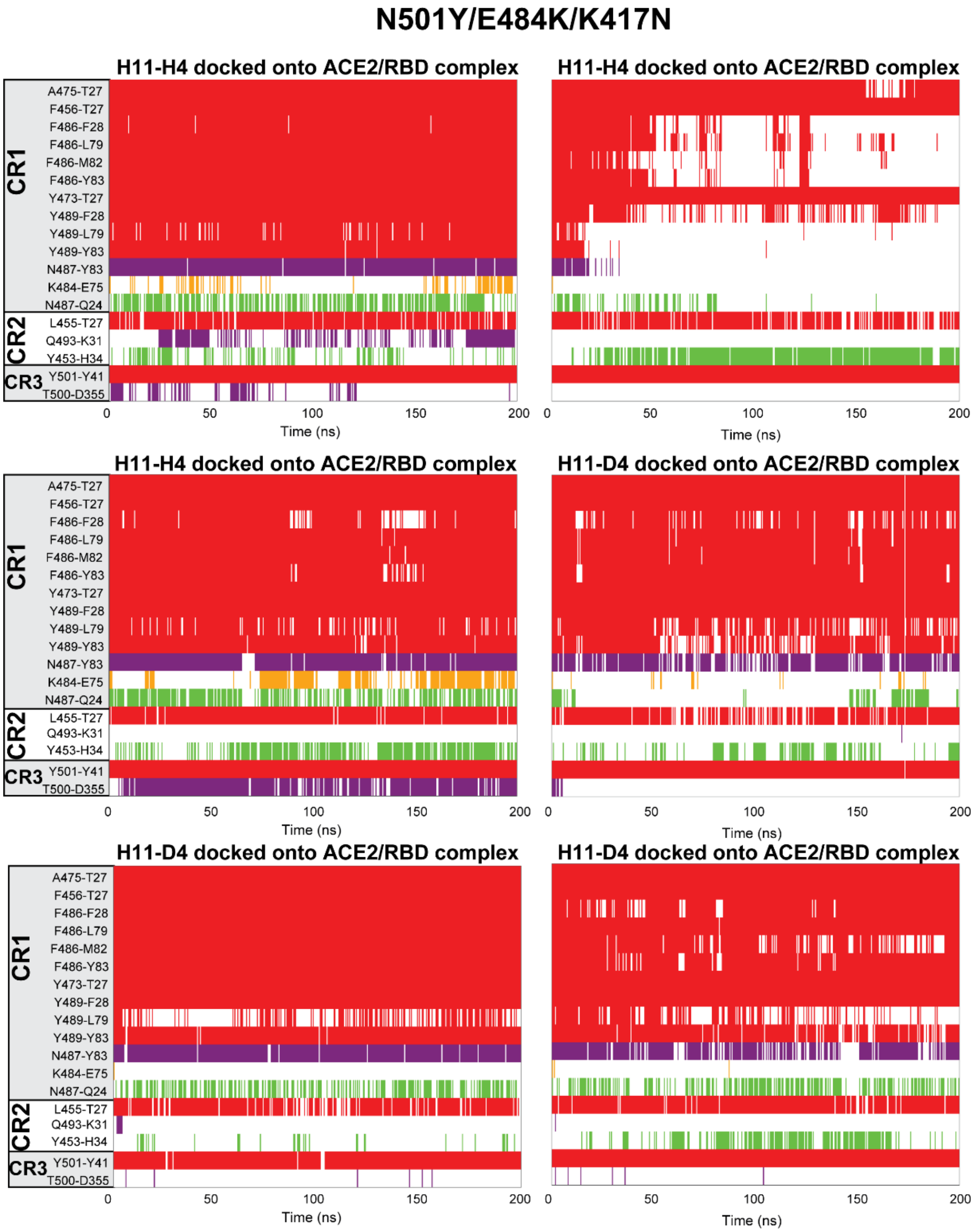


**Figure S10.** Effect of H11-H4 binding on hydrophobic interactions (red), hydrogen bonding (purple), electrostatic interactions (green), and salt bridges (orange) between 501.V2 variant RBD and ACE2**.** Time zero indicates the time instant after minimization.

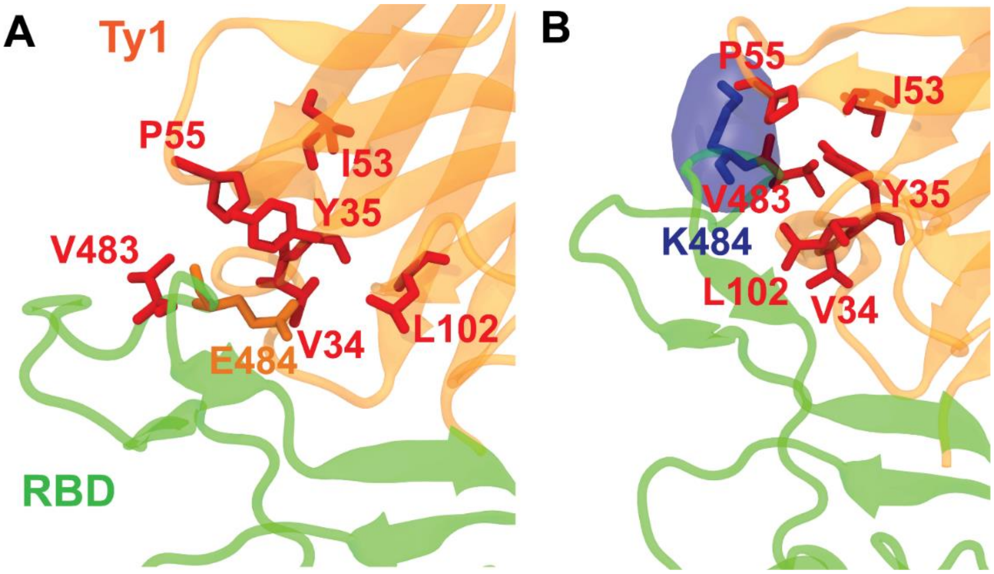


**Figure S11.** Interaction network change of Ty1 and RBD with the N501Y/E484K/K417N mutations. A) Interaction network in WT RBD and Ty1. B) Interaction network in 501.V2 variant and Ty1.

**Table S1.** Starting conformations and durations of the MD simulations performed.

| Run ID | Initial state | RBD sequence | Simulation Type | Simulation Duration (ns) |
|---|---|---|---|---|
| 1a-b | RBD-ACE2 complex (6M0J) | WT SARS-CoV-2 | MD | (a) 100, (b) 100 |
| 1c-d | Final conformers of MD 1a-b | WT SARS-CoV-2 | sSMD | (c) 600, (d) 600 |
| 1e-f | Final conformers of MD 1a-b | WT SARS-CoV-2 | SMD | (e) 200, (f) 200 |
| 2a-b | RBD-H11-H4 complex (6ZBP) | WT SARS-CoV-2 | MD | (a) 200, (b) 200 |
| 2c-d | Final conformers of MD 2a-b | WT-SARS-CoV-2 | sSMD | (c) 600, (d) 600 |
| 2e-f | Final conformers of MD 2a-b | WT-SARS-CoV-2 | SMD | (c) 200, (d) 200 |
| 3a-b | RBD-H11-D4 complex (6YZ5) | WT SARS-CoV-2 | MD | (a) 200, (b) 200 |
| 3c-d | Final conformers of MD 3a-b | WT-SARS-CoV-2 | sSMD | (c) 600, (d) 600 |
| 3e-f | Final conformers of MD 3a-b | WT-SARS-CoV-2 | SMD | (c) 200, (d) 200 |
| 4a-b | RBD-Ty1 complex (6ZXN) | WT SARS-CoV-2 | MD | (a) 200, (b) 200 |
| 4c-d | Final conformers of MD 4a-b | WT-SARS-CoV-2 | sSMD | (c) 600, (d) 600 |
| 4e-f | Final conformers of MD 4a-b | WT-SARS-CoV-2 | SMD | (c) 200, (d) 200 |
| 5a-b | RBD-ACE2 complex (6M0J) | N501Y.V1 variant | MD | (a) 200, (b) 200 |
| 5c-d | Final conformers of MD 5a-b | N501Y.V1 variant | sSMD | (c) 600, (d) 600 |
| 5e-f | Final conformers of MD 5a-b | N501Y.V1 variant | SMD | (e) 200, (f) 200 |
| 6a-b | RBD-H11-H4 complex (6ZBP) | N501Y.V1 variant | MD | (a) 200, (b) 200 |
| 6c-d | Final conformers of MD 6a-b | N501Y.V1 variant | sSMD | (c) 600, (d) 600 |
| 6e-f | Final conformers of MD 6a-b | N501Y.V1 variant | SMD | (c) 200, (d) 200 |
| 7a-b | RBD-H11-D4 complex (6YZ5) | N501Y.V1 variant | MD | (a) 200, (b) 200 |
| 7c-d | Final conformers of MD 7a-b | N501Y.V1 variant | sSMD | (c) 600, (d) 600 |
| 7e-f | Final conformers of MD 7a-b | N501Y.V1 variant | SMD | (c) 200, (d) 200 |
| 8a-b | RBD-Ty1 complex (6ZXN) | N501Y.V1 variant | MD | (a) 200, (b) 200 |
| 8c-d | Final conformers of MD 8a-b | N501Y.V1 variant | sSMD | (c) 600, (d) 600 |
| 8e-f | Final conformers of MD 8a-b | N501Y.V1 variant | SMD | (c) 200, (d) 200 |
| 9a-b | RBD-ACE2 complex (6M0J) | N501.V2 variant | MD | (a) 200, (b) 200 |
| 9c-d | Final conformers of MD 9a-b | N501.V2 variant | sSMD | (c) 600, (d) 600 |
| 9e-f | Final conformers of MD 9a-b | N501.V2 variant | SMD | (e) 200, (f) 200 |
| 10a-b | RBD-H11-H4 complex (6ZBP) | N501.V2 variant | MD | (a) 200, (b) 200 |
| 10c-d | Final conformers of MD 10a-b | N501.V2 variant | sSMD | (c) 600, (d) 600 |
| 10e-f | Final conformers of MD 10a-b | N501.V2 variant | SMD | (c) 200, (d) 200 |
| 11a-b | RBD-H11-D4 complex (6YZ5) | N501.V2 variant | MD | (a) 200, (b) 200 |
| 11c-d | Final conformers of MD 11a-b | N501.V2 variant | sSMD | (c) 600, (d) 600 |
| 11e-f | Final conformers of MD 11a-b | N501.V2 variant | SMD | (c) 200, (d) 200 |
| 12a-b | RBD-Ty1 complex (6ZXN) | N501.V2 variant | MD | (a) 200, (b) 200 |

| | | | | |
|---|---|---|---|---|
| 12c-d | Final conformers of MD 12a-b | N501.V2 variant | sSMD | (c) 600, (d) 600 |
| 12e-f | Final conformers of MD 12a-b | N501.V2 variant | SMD | (c) 200, (d) 200 |
| 13a | RBD-H11-H4+ACE2 complex (6ZBP+(6M0J:ACE2)) | WT SARS-CoV-2 | MD | (a) 200 |
| 14a-c | RBD-ACE2+H11-H4 complex (6M0J+6ZBP:H11-H4) | WT SARS-CoV-2 | MD | (a) 200, (b) 200, (c) 200 |
| 15a-b | RBD-ACE2+H11-H4 complex (Final conformers of MD 1a-b+6ZBP:H11-H4) | WT SARS-CoV-2 | MD | (a) 200, (b) 200 |
| 16a-b | RBD-ACE2+H11-D4 complex (6M0J+6YZ5:H11-D4) | WT SARS-CoV-2 | MD | (a) 200, (b) 200 |
| 17a-c | RBD-ACE2+H11-H4 complex (Final conformers of MD 5a-b+6ZBP:H11-H4) | N501Y.V1 variant | MD | (a) 200, (b) 200, (c) 200 |
| 18a-c | RBD-ACE2+H11-D4 complex (Final conformers of MD 5a-b+6YZ5:H11-D4) | N501Y.V1 variant | MD | (a) 200, (b) 200, (c) 200 |
| 19a-c | RBD-ACE2+H11-H4 complex (Final conformers of MD 9a-b+6ZBP:H11-H4) | N501.V2 variant | MD | (a) 200, (b) 200, (c) 200 |
| 18a-c | RBD-ACE2+H11-D4 complex (Final conformers of MD 9a-b+6YZ5:H11-D4) | N501.V2 variant | MD | (a) 200, (b) 200, (c) 200 |
